# The SAM Domain of EphA2 Inhibits Ligand-Independent Clustering and Activation

**DOI:** 10.1101/2023.10.05.561050

**Authors:** Xiaojun Shi, Ryan Lingerak, Pravesh Shrestha, Matthias Buck, Bing-Cheng Wang, Adam W. Smith

## Abstract

Eph receptors are the largest family of receptor tyrosine kinases (RTKs). They play a role in the pathogenesis of various diseases including cancer, atherosclerosis, fibrosis, infectious diseases, diseases of the central nervous system and age-related cataract. EphA2 has attracted much attention over the years owing to its dysregulation in many diseases. Previous studies have revealed the unique molecular organizations of Eph receptors, and particularly EphA2, into large clusters of receptor-ligand complexes. One unique feature of Eph receptors is a C-terminal sterile alpha motif (SAM) domain, which has been proposed to alter dimerization and kinase activity in EphA2. However, the precise role of the SAM domain in regulating the function and oligomerization state of EphA2 has not been reported. Here we apply a time-resolve fluorescence spectroscopy, PIE-FCCS, to characterize the oligomerization state of EphA2 in live cells and determine the role of the SAM domain. We deleted the SAM domain in the context of full length EphA2 and an intracellular domain (ICD) construct to assess the effect of the SAM domain on oligomerization state, kinase activity, and cellular behavior. Overall, we find that the SAM domain inhibits ligand-independent clustering and kinase activity in both full-length EphA2 and the isolated ICD construct at the cell membrane. These results are consistent with the autoinhibitory features of the C-terminal tail of EGFR and may help resolve the allosteric regulation of other RTKs.

## Introduction

Erythropoietin-producing human hepatocellular (Eph) receptors are the largest family of receptor tyrosine kinases (RTKs). Binding to membrane-bound ephrin ligands, EphA2 forms extensive Eph/ephrin clusters at cell-cell junctions(*1-3*). The formation of such clusters stimulates kinase activity and triggers signaling pathways that affect cellular migration and morphology(*4*). Eph receptors and ephrins play a role in the pathogenesis of various diseases including cancer, atherosclerosis, fibrosis, infectious diseases, diseases of the central nervous system and age-related cataracts (*5, 6*). The overall topology of Eph receptors is similar to other RTKs in that they are type-I transmembrane proteins with an extracellular ligand binding domain and an intracellular kinase domain. The extracellular part of the receptor contains a ligand-binding domain (LBD), a sushi domain, an EGF-like domain and two fibronectin type III (FNIII) domains. The receptor is embedded in the cell membrane via a single alpha-helical transmembrane (TM) domain. The intracellular portion of the receptor consists of a juxtamembrane segment (JMS), a tyrosine kinase domain (KD), a sterile alpha motif (SAM) domain and a PDZ binding motif (*7*). Eph receptors are the only RTKs with a C-terminal SAM domain, and the regulatory function of the SAM domains is still not fully understood.

EphA2 forms extensive signaling arrays upon binding ephrin ligand at cell-cell junctions(*8, 9*). Eph receptors form much larger clusters in the plasma membrane than other RTKS, which was explained by the discovery of multiple binding interfaces (LBD-LBD, sushi-sushi, FNIII1-FNIII1 and LBD-FNIII2) in structural studies of the EphA2 ecto-domain (ECD) (*8, 10, 11*). Based on these findings, a seeding mechanism was proposed, in which Eph/ephrin clusters can recruit unbound EphA2 into extensive signaling platforms(*8, 12*). A recent study carried out a biophysical structural characterization of the intracellular domain (ICD) of EphA2 and substantiated the importance of the SAM domain and its linker in regulating EphA2 signaling (*13*). Our previous study showed that the SAM domain has an inhibitory function on EphA2 kinase activity (*14*). We also used single-color fluorescence correlation spectroscopy (FCS) to reveal a decrease in mobility and increase in molecular brightness of the EphA2 upon SAM deletion. However, the details of how the SAM domain regulates EphA2 kinase activity and oligomerization is still not clear. In this study, we used dual-color pulsed interleaved excitation-fluorescence cross-correlation spectroscopy (PIE-FCCS) to quantify the oligomerization of EphA2 in live cells, with a focus on the SAM domain of EphA2. We also conducted biochemical and cell assays to determine the effect of the SAM domain on EphA2 kinase activity and cell migration.

PIE-FCCS is a time-resolved fluorescence spectroscopy that is well-suited for studying the interaction of membrane proteins in live cell membranes (*15-18*). Using a cross-correlation analysis, the oligomerization state of receptors can be quantified directly (*19*). The results of this work indicate that EphA2 receptors form higher order oligomers upon SAM deletion. We demonstrate that the SAM domain inhibits dimerization of the ICD at the plasma membrane and thus prevents auto-activation of the kinase.

## Results

### PIE-FCCS measures the degree of oligomerization of membrane proteins in live cells

PIE-FCCS utilizes two overlapping laser beams to measure the diffusion of membrane proteins in live cell membranes. The membrane proteins are divided into two populations according to the emission wavelength of their C-terminal fluorescent protein tags. In the PIE-FCCS experiments below, GFP and mCherry are used to label the membrane protein of interest, which allows the diffusion of the “green” (520 nm) and “red” (612 nm) labeled populations to be measured (**Fig. 1A**). The results are reflected in the auto-correlation curves (**Fig. 1B**, green and red curves), which report on the density and mobility of the membrane protein of interest based the amplitude (G_g_(0) and G_r_(0)) and decay time (t_D_) of the curves, respectively. Moreover, the co-diffusion of the green and red population can also be quantified. This provides a tool to measure the oligomerization of the membrane proteins of interests, as the interacting membrane proteins will diffuse together as a complex. The positive amplitude of the cross-correlation curve (**Fig. 1B**, blue curve) is the readout for membrane protein co-diffusion. The cross-correlation value (*f*_*c*_) is calculated using the amplitudes of the auto- and cross-correlation curves to provide a semi-quantitative scale for the degree of oligomerization of the membrane proteins.

**Figure 1.**
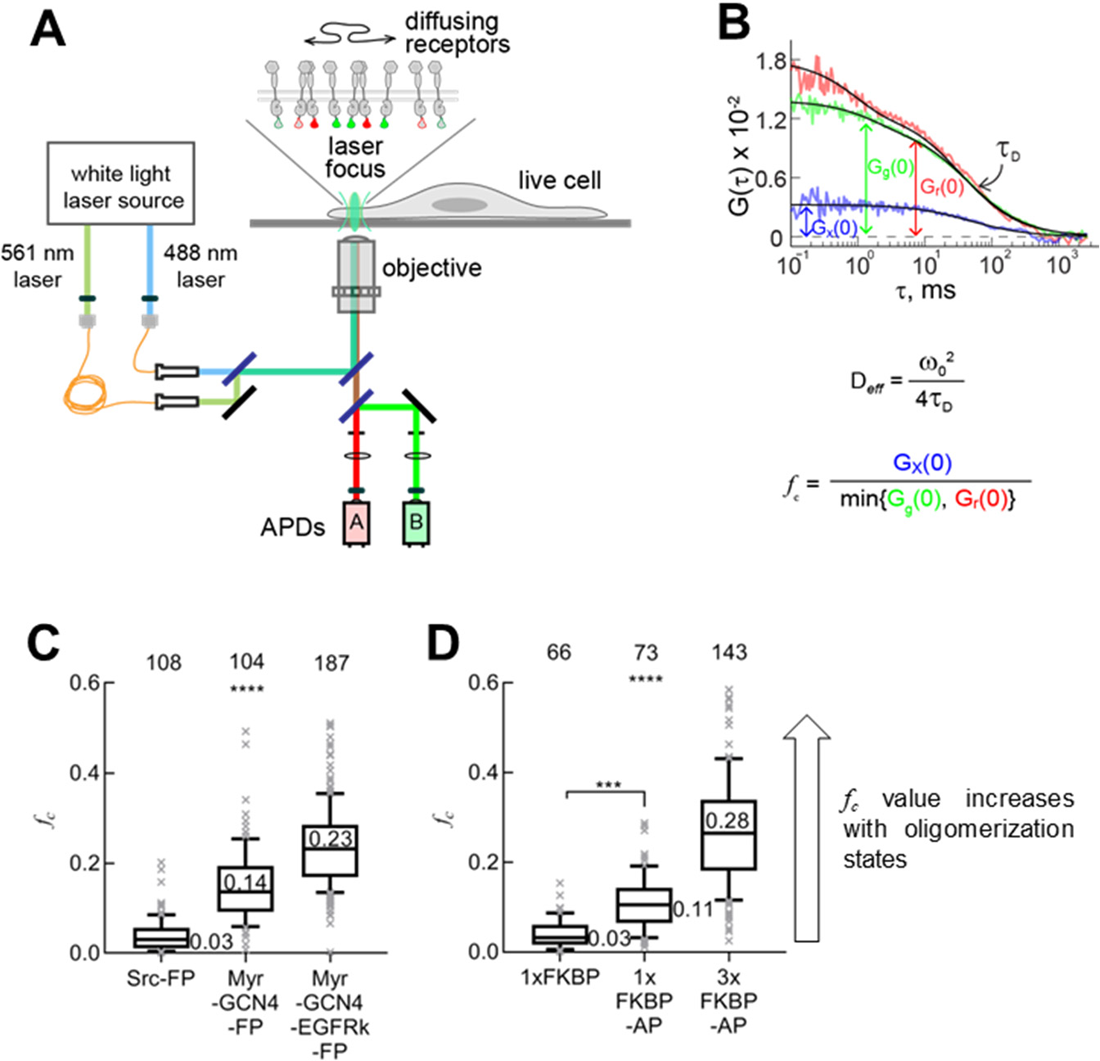
**A)** Schematic diagram of PIE-FCCS experiment. 488 nm and 561 nm pulsed laser beams were focused onto the membrane of cultured COS7 cells expressing fluorescent protein (FP: -GFP and –mCherry) tagged protein of interest. Different lengths of optical fibers were used to create a 50 ns delay between the 488 nm and 561 nm pulses. Single photons in two spectral regions were recorded and time-tagged for analysis. **B)** Representative FCCS curves from measurements on COS7 cells. t_D_ reports on mobility of the diffusive proteins and is used to calculate the effective diffusion coefficients (D_*eff*_). The initial amplitudes of the auto-correlation function curves (green and red curves), G_g_(0) and G_r_(0), and the those of the cross-correlation function curve (blue curve), G_x_(0), is related to *f*_c_ parameter which reports on the multimerization state of the diffusive proteins. **C)** The medians of *f*_c_ values of GCN4 controls. Monomer control: Src-FP (first column) with an *f*_c_ value of 0.03; dimer control: Myr-GCN4-FP (second column) with an *f*_c_ value of 0.14; multimer control: Myr-GCN4-EGFRk-FP (third column) with an *f*_c_ value of 0.23. **D)** The medians of *f*_c_ values of FKBP controls. Monomer control: 1xFKBP (first column) with an *f*_c_ value of 0.03; dimer control: 1xFKBP with AP dimerizer (second column) with an *f*_c_ value of 0.11; multimer control: 3xFKBP with AP dimerizer (third column) with an *f*_c_ value of 0.28. The results of these two control systems indicate that the *f*_c_ value increases with the increase of the oligomerization degree of the membrane protein. These controls have been reported in our previous studies. The *f*_c_ values were presented in box plots with the whiskers showing the 90 percentile of the whole data set. The median values were reported next to the box plots. Each data point was the average of five 10s FCCS measurements performed on one cell. The numbers on top the plots are the total number of cells used. The one-way ANOVA tests were performed to obtain p values (****: p<0.0001, ***: p<0.001).

PIE-FCCS is a sensitive method for measuring membrane protein interactions in live cell membranes. We showed that PIE-FCCS can also distinguish oligomerization from simple dimerization (*20*). A mathematical model was developed to show that the apparent cross-correlation value (*f*_*c*_) increases as the degree of oligomerization increases (*19*). In a system with dimer formation, *f*_*c*_ is greater than 0, but does not reach the ideal value of 1 even for complete dimerization, because the formation of same-color dimers (GFP-GFP or mCherry-mCherry) are “invisible” to cross-correlation analysis (*19*). This trend is verified with two sets of control proteins that we developed for PIE-FCCS experiments. There is a monomer control, a dimer control and an oligomer control in each set of proteins. The details of these two sets of control proteins can be found in our previous works (*19, 20*). The results of the two sets of controls are summarized in **Fig. 1C**. It is clear in both sets of data that the *f*_*c*_ value increases with the degree of oligomerization. In addition to the *f*_c_ value, the mobility of membrane proteins can also be obtained from PIE-FCCS measurements. Comparing the mobility of proteins with different oligomerization states corroborates our interpretation of the cross-correlation analysis, as larger oligomers will exhibit slower diffusion.

### The SAM domain prevents EphA2 clustering prior to ligand stimulation

We carried out dual-color PIE-FCCS experiments to assess the effect of SAM domain on the oligomerization of EphA2. The experiments were carried in COS7 cells that are co-transfected with GFP and mCherry tagged EphA2 with or without the SAM domain (**Fig. 2A**). The relative change in the extent of oligomerization of these protomers can be assessed using the change of *f*_c_ values. First, we stimulated EphA2WT expressing cells with ephrinA1-Fc (EA1-Fc) ligand. We observed a dramatic increase of the *f*_c_ value from 0.21 to 0.52 upon EA1-Fc stimulation of EphA2WT (**Fig. 2B**). This indicates a dramatic increase in the oligomerization of EphA2 in response to ligand stimulation, which agrees with the observation of ligand-induced clustering of EphA2 reported in previous studies (*8, 10*). The *f*_c_ value of the SAM deletion mutant (EphA2DS) without ligand is 0.45 (**Fig. 2B**, third column). Comparing to the *f*_c_ value of EphA2WT (0.24, **Fig. 2B**, first column), the increase in *f*_c_ value of EphA2DS indicates the increase in oligomerization is caused by SAM deletion. In fact, the *f*_c_ value of EphA2DS is similar to that of the EA1-Fc stimulated EphA2WT (0.52, **Fig. 2B**, second column), indicating that SAM deletion causes clustering of EphA2 similar to the ligand-induced EphA2 clustering.

**Figure 2.**
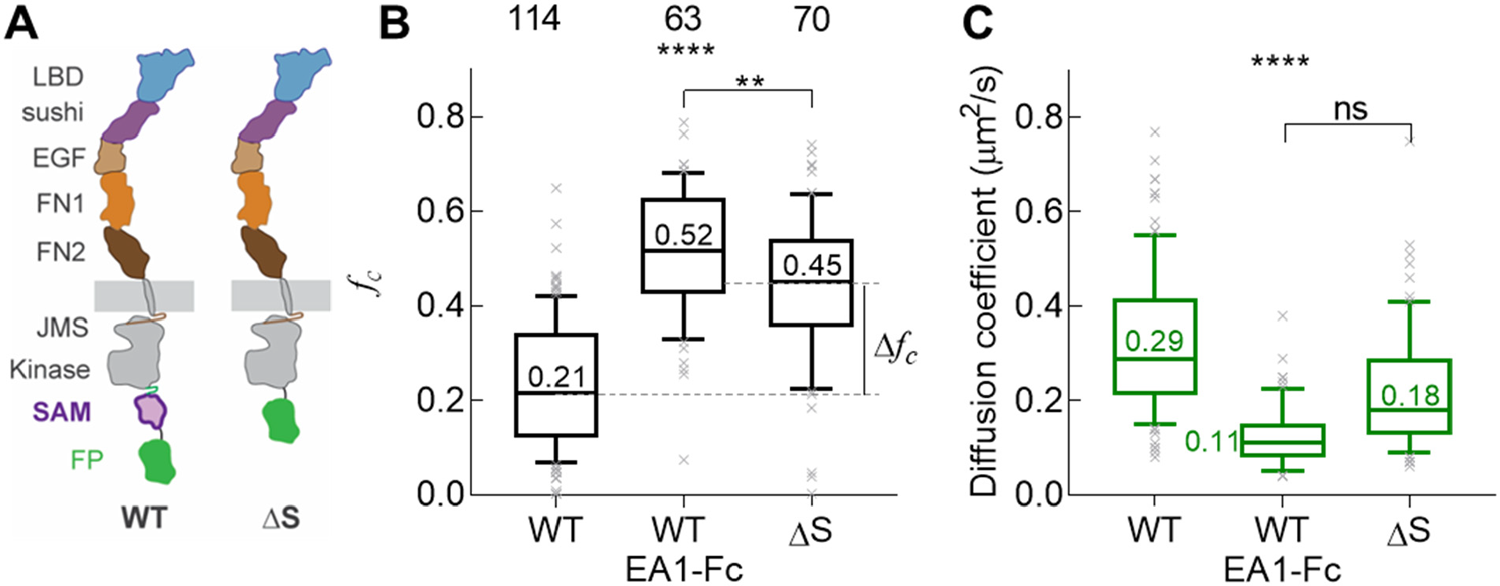
**A)** Schematic diagram of EphA2WT and EphA2ΔS. **B)** *f*_c_ values of EphA2WT without and with EA1-Fc ligand stimulation, and EphA2ΔS. The *f*_c_ value of EphA2ΔS (0.45) is larger than EphA2WT (0.24) and similar to EA1-Fc stimulated EphA2WT (0.52), indicating EphA2ΔS form clusters similar to EA1-Fc stimulated EphA2WT. **C)** Diffusion coefficients of EphA2WT without and with EA1-Fc ligand stimulation, and EphA2ΔS. Data were presented in box plots with the whiskers showing the 90 percentile of the whole data set. The median values were reported next to the box plots. The numbers on top the plots are the total number of cells used. One-way ANOVA tests were performed to obtain the p values (****: p<0.0001, **: p<0.01, ns: not significant).

The mobility of these EphA2 constructs was obtained from the same set of PIE-FCCS measurements and is summarized in **Fig. 2C**. EphA2ΔS shows slower diffusion compared to EphA2WT, which supports the clustering of EphA2 due to deletion of the SAM domain. All these results suggest that the SAM domain prevents EphA2 from clustering prior to ligand stimulation. Clustering of EphA2 is a significant molecular event because it is coupled to the activation of kinase activity. In our previous study (*14*), we found an elevated phosphorylation in the di-tyrosine motif of the JMS upon EphA2 SAM deletion, which indicated the increased activation of the EphA2 kinase. This observation suggests that the SAM domain is involved in the regulation of EphA2 kinase activity through preventing the ligand-independent clustering of the receptor. In order to gain more details of the regulatory function of SAM domain of EphA2, we turned to a study of the intracellular domain (ICD) of EphA2.

### The SAM domain inhibits interactions between the EphA2 kinase domains

We next deleted the TM and ECD domains from EphA2 and conducted PIE-FCCS measurements of these truncated constructs. To anchor the construct to the plasma membrane, we fused it to a membrane localization sequence used in our previous studies (N-terminal myristoylation of a c-Src-derived peptide) to create Myr-ICD (**Fig. 3A**). The SAM domain truncation mutant was created from the Myr-ICD sequence to yield Myr-ICD-ΔS (**Fig. 3A**). This construct was transiently transfected to COS7 cells to perform PIE-FCCS measurements. The results indicate that the ICD of EphA2 (Myr-ICD) is monomeric, as its *f*_c_ value is close to 0 (**Fig. 3B**). The interpretation of this observation is that the presence of the SAM domain inhibits any observable clustering, which is surprising given that the canonical function of SAM domains is to promote dimerization and oligomerization (*21*). However, our previous in vitro studies have shown that the EphA2 SAM domain does not form homodimers or oligomers up to high protein concentrations (*22*). Similar to the full-length receptor, deletion of the SAM domain from Myr-ICD leads to increased oligomerization, demonstrated by the median *f*_c_ value of Myr-ICD-DS of 0.14 (**Fig. 3B**). This *f*_c_ value is consistent with dimerization, as it is similar to a dimer control (**Fig.1C&D**). Moreover, the mobility of Myr-ICD-ΔS is significantly lower than that of Myr-ICD (**Fig. 3C**), which agrees with the formation of dimers upon SAM deletion. Together, our observations show that the EphA2 kinase domain by itself has a strong propensity for dimerization that is inhibited by the SAM domain.

**Figure 3.**
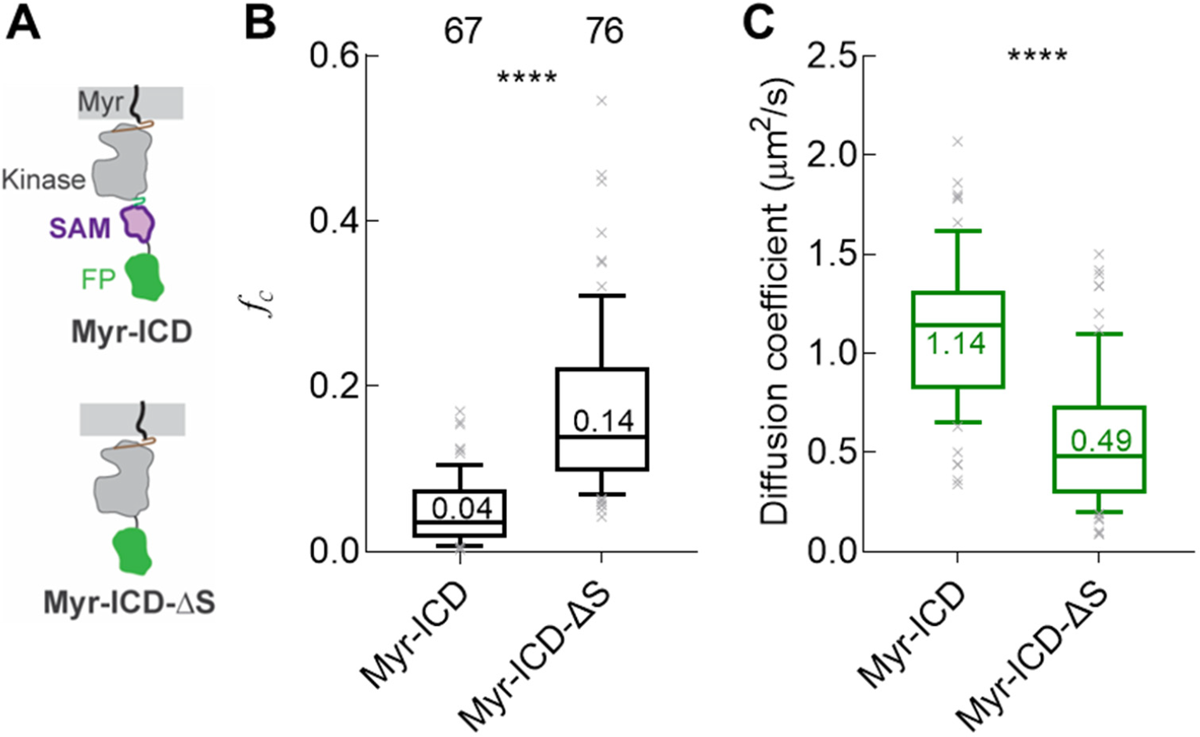
**A)** Schematic diagram of isolated EphA2 intracellular domain (ICD), Myr-ICD and Myr-ICD-ΔS. **B)** *f*_c_ values of Myr-ICD and Myr-ICD-ΔS. The *f*_c_ value of Myr-ICD-ΔS (0.14) is larger than Myr-ICD (0.04), indicating that Myr-ICD-ΔS undergo dimerization. **C)** Diffusion coefficients of Myr-ICD and Myr-ICD-ΔS. Two-tailed T tests were performed to obtain the p values (****: p<0.0001).

### Intrinsic activity of the isolated EphA2 intracellular domain (ICD)

The kinase domains of EGFR, FGFR1 and FGFR2 form asymmetric dimers to enable trans-phosphorylation and activation (*23-25*). We hypothesize that the dimerization of EphA2 kinase domains we observed above also enables trans-phosphorylation and activation of EphA2. An antibody against the phospho-dityrosine motif conserved in Eph receptors (pY-EphA/B) was used to detect the extent of phosphorylation the of EphA2 kinase, a consequence of its own kinase activity, in a Western Blot analysis. Upon SAM deletion, the pY-EphA/B level was increased (**Fig. 4A**), indicating trans-phosphorylation of the kinase. In an immunofluorescence analysis, transfected cells were fixed and permeabilized to allow antibody-staining. The pY772 antibody, which is against the phospho-tyrosine (772) on the activation loop of the kinase, was used as primary antibody, and a secondary antibody (anti-rabbit AlexaFluor 566) was used to visualize the phosphorylation level. The relative phosphorylation level of Y722 was indicated with the ratio of the intensity of the GFP channel and the pY772 channel, which represents the expression level of the protein and its level of phosphorylation, respectively (**Fig. 4B**). The images show that the pY772/GFP ratio at the plasma membrane increased due to SAM deletion (**Fig. 4C**), which suggests elevated phosphorylation levels at Y772. Together, these results indicate that the dimerization of the kinases induced by SAM deletion leads to elevated phosphorylation of several key tyrosine motifs suggesting activation of the kinase.

**Figure 4.**
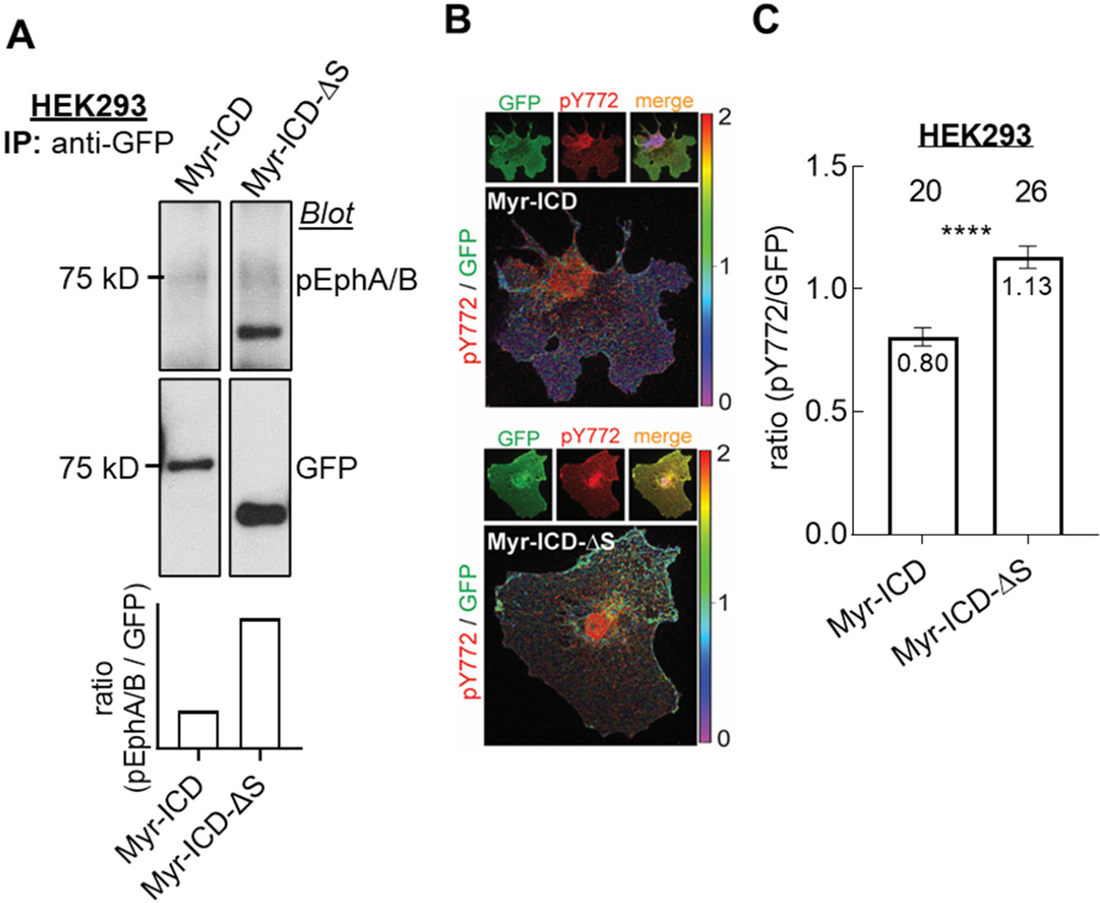
**A)** Total cell lysates from HEK293 cells expressing Myr-ICD and Myr-ICD-ΔS were immunoblotted with the indicated antibodies. The pY-EphA/B antibody was raised against the phosphorylated di-tyrosine motif in the JMS conserved in both EphA and EphB receptors. Total expression levels were detected with an antibody against the GFP tag. **B)** Confocal images of HEK293 with expression of the GFP-tagged Myr-ICD (top) and Myr-ICD-ΔS882 (bottom). The cells were fixed and stained with pY772 antibody (rabbit) and then visualized with anti-rabbit AlexFlour 566 to indicate the phosphorylation level of Y772 (pY772). The images of GFP-tagged protein (GFP), anti-rabbit AlexFlour 566 (pY772) and the merged image with DAPI nuclei staining are shown on the top panels. The ratio of the intensity of pY772 to the intensity of GFP (pY772/GFP) is mapped for the cell images to indicate the different strength of kinase activity of both constructs. **C)** Statistics of the pY772/GFP ratio of cells expressing Myr-ICD and Myr-ICD-ΔS. The average values are reported in the bars. The error bars represent the SEM. The numbers on the top indicate the total numbers of cells used in the statistics. Two-tailed T test was performed to obtain the p value (****: p<0.0001).

We also performed migration experiments on HEK293 cells with stable expression of Myr-ICD and Myr-ICD-ΔS. The results indicate that Myr-ICD-ΔS expressing cells move slower than those expressing Myr-ICD (**Fig. 5**). As activation of EphA2 kinase inhibits cell motility (*26*), the results of the wound-healing assay support the conclusion that the kinase domain of Myr-ICD-ΔS is activated through dimerization. These measurements on the membrane bound, but extracellular region-trunctated EphA2, EphA2-ICD, suggest that the SAM domain inhibits kinase-kinase interaction, and once this inhibitory mechanism is removed – in this case by deletion of the SAM domain - the kinases can undergo dimerization which leads to the increase of kinase activity.

**Figure 5.**
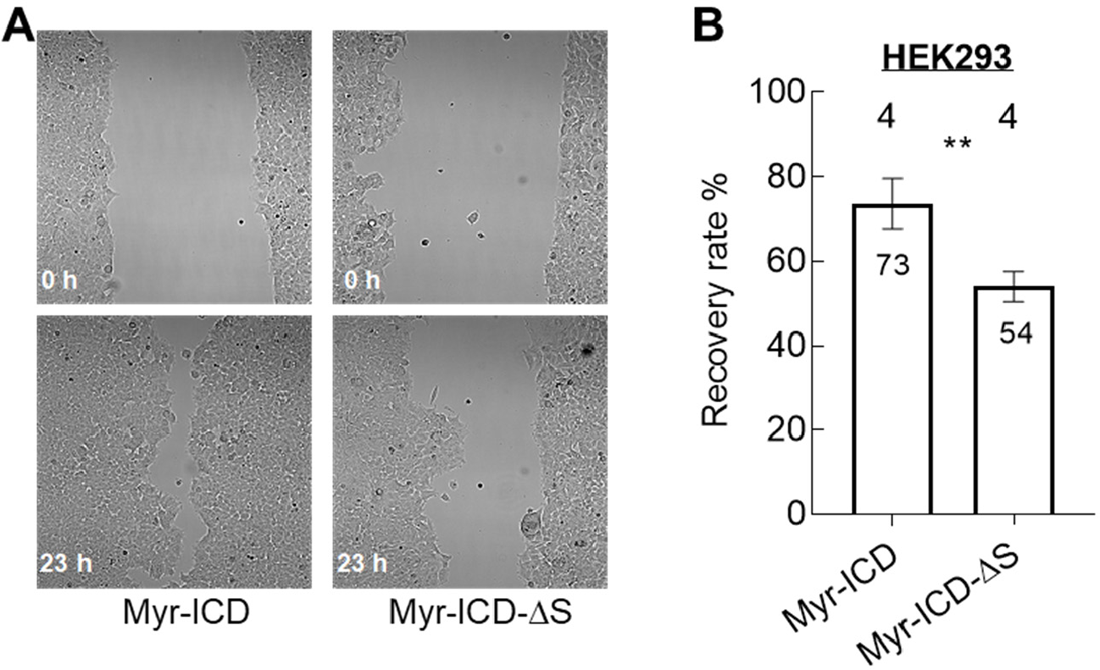
**A)** Sample bright field images of wounds at 0 h and 23 h of Myr-ICD and Myr-ICD-ΔS expressing HEK293 cells. **B)** Statistics of the wound recovery rate of cells expressing Myr-ICD and Myr-ICD-ΔS. The average values are reported in the bars. The error bars represent the SEM. A total of 4 sets of wound were used for each construct. Two-tailed T test was performed to obtain the p value (**: p<0.01).

## Discussion

In this study, we showed that PIE-FCCS can be used to monitor changes in oligomerization for membrane proteins in live cell membranes. Specifically, the cross-correlation value (*f*_c_) correlates with the extent of oligomerization. We observed a dramatic increase of *f*_c_ value, from 0.21 to 0.52 (**Fig. 2B**), of EphA2 in response to ligand stimulation. This shows that EphA2 undergoes clustering in response to ligand, which is consistent with previous studies using different methods (*27-29*). Previously, deletion of the SAM domain was reported to cause increased kinase activity in EphA2 (*14*). The underlying mechanism of this phenomenon has been previously studied by a FRET assay and by us using single-color FCS (*14, 27*). The FRET assay concluded that the deletion of SAM domain stabilizes the EphA2 dimer by increasing the dimerization affinity. However, the results of PIE-FCCS measurements in this study clearly indicate that the degree of EphA2 oligomerization increases upon deletion of SAM domain, as we observed an increase in *f*_c_ value from 0.21 to 0.45 (**Fig. 2B**). In fact, the *f*_c_ values of ligand stimulated EphA2 and EphA2 with SAM deletion are so similar (**Fig. 2B**) that we can conclude that SAM deletion causes EphA2 to cluster to the similar level as it is caused by ligand stimulation. Such clustering of EphA2 upon ligand stimulation has been reported to be a hallmark of EphA2 RTK activity (*10*). Paradoxically, the SAM domain is generally reported to facilitate protein-protein interactions (*21, 30*), but the EphA2 SAM does not form dimers or homo-oligomers in solution (*22*). By contrast, the SAM domain of EphA3 was reported to promote the dimerization of EphA3 (*31*). Thus, in the case of EphA2, the SAM domain performs the opposite function by acting as an inhibitor to clustering of EphA2.

We further investigated the behavior of the isolated ICD of EphA2 to provide more details of the unique function of the SAM domain of EphA2. From our data we conclude that the ICD of EphA2 (including JMS, kinase and SAM domain) is mostly monomeric, while deletion of SAM domain from the isolated ICD causes dimerization. Our biochemical study of the isolated ICD shows that SAM deletion causes increased phosphorylation levels of a key activation loop tyrosine in the ICD. This suggests the activation of the EphA2 kinase and is supported by a cell migration assay. The canonical model of RTK activation describes the dimerization of kinase domains to facilitate trans-phosphorylation of tyrosine motifs. The behavior of isolated ICD of EphA2 fits this model, as the activity of kinase increases upon dimerization caused by the deletion of SAM domain from the monomeric full-length ICD. This also suggests that EphA2 kinase has a strong propensity for dimerization. Similarly, kinases of other RTKs, such as EGFR, FGFR1 and FGFR2, form dimers that regulate kinase activity (*23-25*). The C-terminal tail of EGFR, which is in the same position as the SAM domain of EphA2, plays an important role in inhibiting the formation of active asymmetric form of EGFR kinases (*32*). In fact, an oncogenic mutant of EGFR that harbors the deletion of C-terminus tail was reported to undergo ligand-independent activation (*33*). In a similar way, the SAM domain of EphA2 might interact with the rest of the ICD as an inhibitory mechanism to prevent unregulated dimerization and activation of the kinase. Deletion of the SAM domain facilitated the formation of active dimers through kinase-kinase interactions. However, the structural details of this mechanism still need to be investigated further.

## Methods

### Plasmids and cell cultures

Full-length EphA2 genes (EphA2FL AA 1-971, EphA2ΔS AA 1-903) were sub-cloned into Gateway donor vectors which were shuttled into pLenti-CMV-Puro vectors for mammalian expression. Intracellular domain of EphA2 genes (Myr-ICD AA 559-971, Myr-ICD-ΔS AA 559-882) were sub-cloned into N1 vectors for mammalian expression. COS7 and HEK293 cells were cultured in DMEM (10% FBS). The culture dishes were coated with collagen for HEK293 cells.

### PIE-FCCS instrumentation and data analysis

PIE-FCCS measurements were performed on a customized Nikon Eclipse Ti inverted microscope (Nikon Corp., Tokyo, Japan) described previously (*18*). A continuum white light laser (9.7 MHz, SuperK NKT Photonics, Birkerod, Denmark) was used as excitation laser source. The source had an internal pulse picker that set the pulse rate to 10 MHz. A wavelength splitter inside the emission box picked off 488 and 561 nm beams that were passed through narrow-band excitation filters (488: LL01-488-12.5; 561: LL02-561-12.5, respectively; Semrock, Rochester, NY) before being coupled into single-mode optical fibers (488: QPMJ-3AF3U-488-3.5/125-3AS-18-1-SP; 561: QPMJ-3AF3U-488-3.5/125-3AS-3-1-SP; OZ Optics, Ottawa, Ontario). The 488 nm beam passed through a 3 m fiber while the 561 nm beam passed through an 18 m fiber. The 15 m-length difference of the two fibers introduced a 50 ns delay between the two pulse trains for pulsed interleaved excitation (PIE). The beams exited the fibers and were collimated with infinity corrected objective lenses (L-10x, Newport, Irvine, CA). Continuously variable ND filters were placed after the lenses to adjust the laser power for each beam. The two beams were overlapped with a 503 nm cutoff dichroic beamsplitter (LM01-503-25, Semrock, Rochester, NY) before being sent into the optical path of the microscope. A customized TIRF filter cube (91032, Chroma Technology Corp., Bellows Falls, VT) with a two-color dichroic mirror and laser blocking filer (zt488/561rpc and zet488/561m, Chroma Technology Corp., Bellows Falls, VT) was used to couple the beams to the objective. A 100X TIRF (oil) objective, NA 1.49, (Nikon Corp., Tokyo, Japan) was used to focus the excitation beams on the sample and collect the emitted photons. The emitted photons passed through a 50 μm pinhole placed at the output port of the microscope to achieve confocal detection. The beam was collimated with a 100 mm focal length achromatic lens (AC254-100-A-ML, Thorlabs Inc., Newton, NJ) and then split into two beams with a 560 nm long-pass beamsplitter (FF560-FDi01-25X36, Semrock, Rochester, NY). The beams were directed through a 520/44 nm emission filter (FF01-520/44-25, Semrock, Rochester, NY) and a 612/69 nm emission filter (FF01-621/69-25, Semrock, Rochester, NY) to obtain the green (520 nm) and red (612 nm) fluorescence signal, respectively. to the signal was collected by single photon avalanche diode (SPAD) detectors (Micro Photon Devices, Bolzano, Italy) with a time-resolution of 30 ps, a 50 μm_2_ active area, and 25 dark counts per second. The photon counts were recorded by a four-channel-routed time-correlated single photon counting (TCSPC) card (Picoharp 300, PicoQuant, Berlin, Germany) synchronized with the white light laser source. Data was processed and analyzed with a home-written Matlab script.

To prepare for PIE-FCCS measurements, cells were plated on MatTek glass bottom culture dish 2-3 days prior to imaging and then transfected 24 hours prior to imaging. Measurements were performed on single cells in Opi-MEM at 37 °C. For ligand stimulation, cells were incubated with 1 μg/mL ligand for 20 min prior to measurements. The laser beams were set at 300 nW (488 nm) and 800 nW (561 nm) and were focused on flat membrane areas near the edge of the cell. Five 10 s measurements were taken on each cell, were fit, and analyzed as described previously (*18, 19*).

### Immune blotting

HEK293 cells were transfected with ICD constructs using Lipofectamine 2000 (Thermo Fisher) and were selected with G418. Cells with stable expression were cultured until subconfluent. Cells were lysed for 30 min at 4 _o_C in modified RIPA buffer (20 mM Tris, pH 7.4, 120 mM NaCl, 1% Triton X-100, 0.5% sodium deoxycholate, 0.1% SDS, 10% glycerol, 5 mM EDTA, 50 mM NaF, 0.5 mM Na_3_VO_4_, and protease inhibitors). Lysates were spun down at 13,000 g for 5 min. Immunoblot was carried out as described (*26*). Antibodies used include mouse anti-GFP, rabbit anti-phospho-EphA/B which was raised against the phosphorylated di-tyrosine motif in the JMS.

### Immune staining

COS7 cells were transfected with ICD constructs using Lipofectamine 2000 and were selected with G418. Cells were treated with 5% paraformaldehyde at 4 _o_C for 15 min and washed with PBS. Fixed cells were permeabilized with 2% Triton X-100 solution and washed with PBS. Then DEME was used for blocking for 1 h. After blocking, cells were stained with primary antibodies (anti-pY772, rabbit, Cell Signaling Technology #8244) for 1 h and then with secondary antibodies (anti-rabbit AlexaFluor 566) for 30 min. After antibody staining, cells were mounted and subjected to confocal fluorescence imaging. All cell samples were imaged under the same illumination condition.

### Cell migration

HEK293 cells were transfected with ICD constructs using lipofectamine2000 and were selected with G418. Cells with stable expression were cultured in m-Dish with 4 micro-well insert (ibidi) until reaches total confluency. Then the block in the plate was removed to create gaps between cell cultures. The gaps were imaged with bright field microscopy at 0 h and after 23 h incubation. The area of the gaps was measurement with ImageJ.

## Acknowledgements

XS was supported by NCI training grant 5T32CA059366. MB was supported by R01GM112491 and R01EY029169. BW was supported by NIH R01CA096533, R01NS096956, and R01CA250067, and also by CCSG CA043703. AWS was supported by the National Science Foundation (CHE-1753060).

